# FastReseg: using transcript locations to refine image-based cell segmentation results in spatial transcriptomics

**DOI:** 10.1101/2024.12.05.627051

**Authors:** Lidan Wu, Joseph M. Beechem, Patrick Danaher

**Affiliations:** Bruker Spatial Biology, Seattle, WA, USA 98105

## Abstract

Spatial transcriptomics (ST) is a rapidly advancing field, yet it is challenged by persistent issues with cell segmentation accuracy, which can bias biological interpretations by making cells appear more similar to their neighbors than they truly are. FastReseg introduces a novel class of algorithm that employs transcriptomic data not to redefine cell boundaries but to rectify inaccuracies within existing image-based segmentation outputs. By combining the rich information from image-based methods with the 3D precision of transcriptomic analysis, FastReseg enhances cell segmentation accuracy. A key innovation of FastReseg approach is its transcript scoring system, which scores each transcript for its goodness-of-fit within host cell using log-likelihood ratio. This scoring system facilitates the quick identification and correction of spatial doublets, *i.e.* cells erroneously segmented due to close proximity or spatial overlap in 2D. FastReseg approach offers several advantages: it reduces the risks of circularity in deriving cell boundaries from expression data and minimizes spatial-dependent biases arising from erroneous segmentation. It also addresses computational challenges often associated with existing transcript-based methods by introducing a heuristic, modular workflow that efficiently processes large datasets, a critical feature given the increasing size of spatial transcriptomics datasets. Its modular workflow allows for individual components to be optimized and seamlessly integrated back into the overall pipeline, accommodating ongoing advancements in segmentation technology. By enabling efficient management of large datasets and providing a scalable solution for refining cell segmentation, FastReseg is poised to enhance the quality and interpretability of spatial transcriptomics data even as underlying image-based cell segmentation techniques evolve.

## Introduction

Cell segmentation is foundational to spatial transcriptomics, as it delineates individual cells and assigns transcript molecules to their cells of origin. The accuracy of segmentation is a critical determinant of data quality in spatial transcriptomics, influencing all subsequent analyses. Segmentation errors can contaminate single-cell expression profiles with transcripts from neighboring cells, introducing spatially dependent bias to the data and confounding analyses. For example, differential expression analyses of a given cell type across spatial domains may be skewed by errors that arise from poorly segmented cells, overshadowing genuine biological signals [1].

To date, most spatial transcriptomics studies employ image-based cell segmentation, and typically involve immunofluorescence (IF) staining of samples with cellular morphological markers such as the nucleus and membrane, allowing for the visual identification of cell boundaries. Methods like the watershed algorithm [2] and nucleus expansion have been widely used in the past, but they have largely been superseded by more advanced techniques that leverage deep learning, such as Cellpose [3] and Mesmer [4]. However, fundamental limits in stained images of real-world tissue samples (**Figure 1**B) make image-based segmentation struggle with issues like weak staining, ambiguous boundaries, and inability to optically resolve vertical overlap of cells, leading to errors that may propagate through subsequent analytical stages.

**Figure 1.**
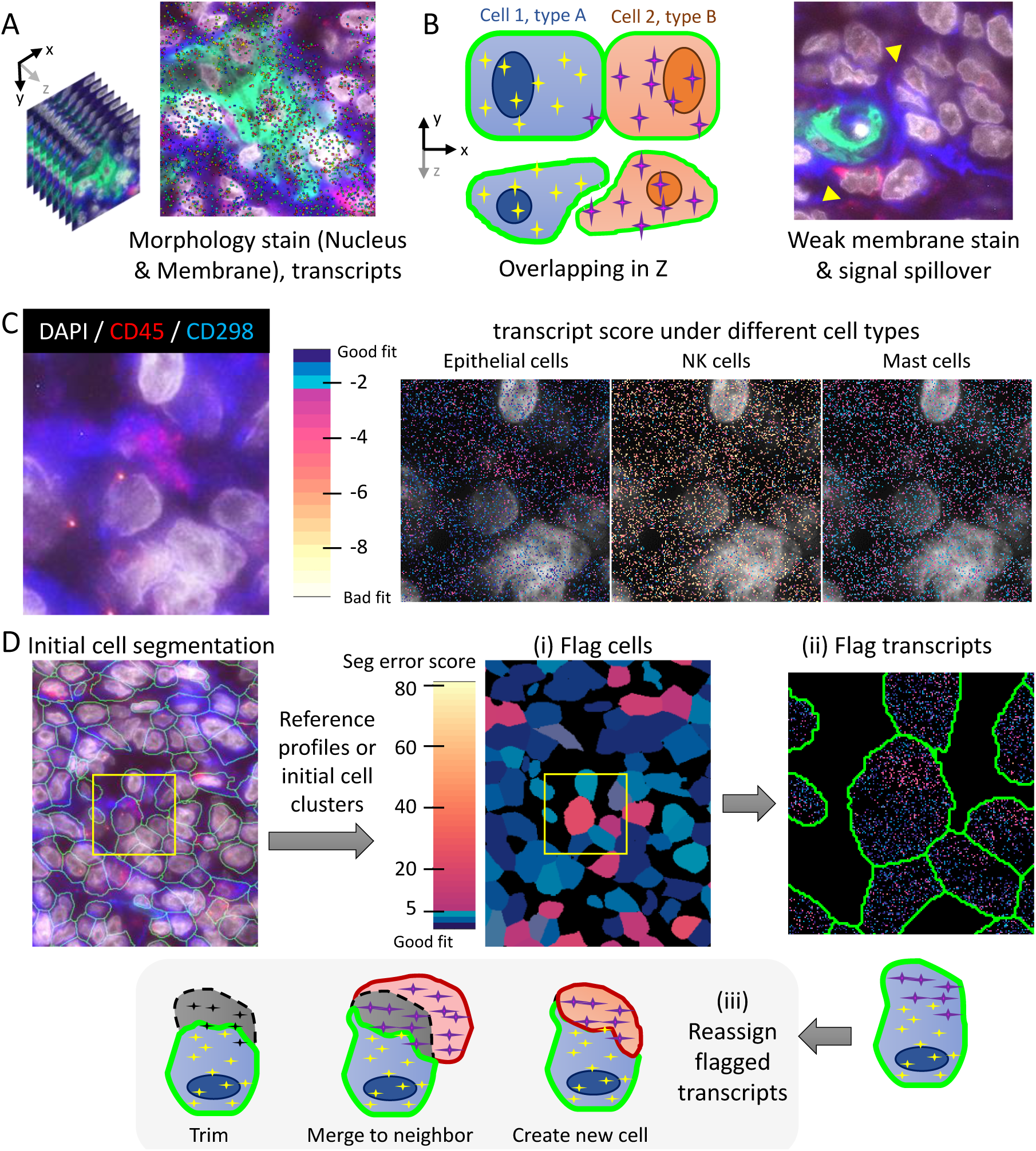
FastReseg, an algorithm for improved cell segmentation in spatial transcriptomics. (A) Visualization of typical spatial transcriptomics data showing morphology stains for nucleus and membrane alongside transcript distribution (shown as dots colored by gene identities), providing essential context for segmentation. (B) Illustration of inherent challenges in image-based cell segmentation, including limited resolution to resolve overlapping cells in the z-axis, weak or incomplete membrane stain in real-world tissue samples (highlighted by yellow arrows), and lateral spillage of optical signals in densely packed tissues. The example images in (A) & (B) are from a melanoma dataset, where antibodies against different protein markers (CD298: Blue, PanCK: Green, CD45: Red) and chemical nuclear stain (DAPI: gray) were used to visualize the sample morphology. (C) Spatial pattern of transcriptional scores (right) based on each gene’s log-likelihood expression under various reference cell types (Epithelial cells, NK cells, Mast cells) could be utilized to identify the presence of different cell types in a spatial context as validated by the orthogonal antibody staining (left, CD298: blue, CD45: red, DAPI: gray). Different color hues and intensities of the dots indicate the goodness of fit of each transcript (dot) under each reference cell type as shown in the color bar, ranging from good (blue) to poor (red and yellow). See definition of tLLR score in Methods section for how to calculate the transcriptional scores plotted here. (D) Overview of the FastReseg workflow. Starting with initial cell segmentation and cluster-specific reference expression profiles, FastReseg scores each transcript based on its goodness-of-fit with respect to most probable cell type given the expression profiles of corresponding host cells and then flags cells with high spatial dependency in the spatial pattern for their transcript scores as cells with putative segmentation errors (i). Within the flagged cells, FastReseg can further identify the transcripts with poor fit, segregate them into spatially distinct groups and flag them as contaminating transcripts from different neighboring cells (ii). In the correction phase (iii), FastReseg evaluates both the expressional and physical spatial context of each flagged transcript group and decides its refinement action where the flagged transcripts are either trimmed to extracellular space, merged with neighboring cells, or reassigned to newly created cells based on a set of heuristic rules, aiming to resolve cell segmentation errors and improve transcript assignment accuracy. See Figure 4 and Methods section for the detailed process.

Transcript-based methods provide an alternative means to segment regions of distinct local expression profiles using the spatial locations and identities of RNA transcripts [5,6]. But they can face difficulties at the borders of closely related cell types or in regions with sparse transcript data, limiting their usage in many cell-driven biological studies. Baysor [7], JSTA [8] and Proseg [9] represent a new class of hybrid approach, taking image-based segmentation as a prior which would be updated using transcript data. Nevertheless, in practice the prior usage is often limited to the nuclear segmentation results and the transcriptomic data frequently dominates the final segmentation outcome, introducing circular reasoning into analyses as the gene expression used for defining cellular boundaries would inadvertently manifest itself in the resulting single-cell expression profiles as segmentation outcomes. Moreover, the existing hybrid methods often require substantial computational resources [7,8,9,10] due to the volume of transcript data and sophisticated modeling involved and thus struggle to generate satisfactory cell-level segmentation for densely packed tissue samples.

To address these challenges, we introduce FastReseg, an R package designed to enhance the precision of cell segmentation by leveraging transcriptomic data to correct and refine initial image-based segmentation results. FastReseg processes spatial transcriptomic datasets, along with their initial cell assignment provided by image-based cell segmentation, through a three-tiered modular approach: it first assesses cells for potential segmentation inaccuracies, then identifies misassigned transcript groups within those likely erroneous cells, and finally reassigns these mislocated transcripts to their appropriate cellular origins. This stepwise progression allows users the flexibility to halt the process after any stage to examine intermediate results, adapting the workflow to suit specific research needs at various analytical depths. The modular design of FastReseg also simplifies the optimization of parameters for each module, facilitating the adaptation of this tool in diverse research contexts.

FastReseg’s benefits are multifold: it reduces the risk of circularity in data analysis by maintaining image data as the primary source for initial segmentation while utilizing transcript data for refinement. This strategy ensures that gene expression data used to study cell functions downstream is not compromised by the very methods intended to define cell boundaries. Besides, as image-based segmentation methods continue to evolve, FastReseg remains a relevant tool, complementing these advancements with its additional layer of transcript-based refinement. Moreover, FastReseg is designed to be fast and memory-efficient, handling computations on a cell-by-cell basis which allows saving the most expensive computations for the cells flagged as high likelihood of segmentation error. Recognizing the three-dimensional nature of tissue structures, FastReseg considers transcripts’ full 3D positions during modeling and is capable of ingesting 3D segmentation results, making it particularly suited to modern, high-resolution spatial transcriptomics datasets.

Through FastReseg, we offer a novel and practical solution to one of the most pressing challenges in the field of spatial transcriptomics, propose a framework that could significantly improve the accuracy and reliability of cell segmentation and thereby enhance the overall quality of biological insights derived from spatial data.

## Results

### Concept of Transcript scoring

FastReseg works on the premise that each RNA transcript offers evidence supporting or refuting the presence of a particular cell type at its location. For example, a CD68 transcript suggests the presence of a macrophage, while simultaneously arguing against the cell being a T cell. This foundational idea is encapsulated by deriving a log-likelihood ratio (*tLLR*) for each gene under each expected cell type against the most probable cell type of given gene across the reference expression profiles for the dataset of interest. These log-likelihood ratios, which we term tLLR scores, are used to score every transcript in every cell in the dataset and serve as the foundation of FastReseg’s operations. While one could obtain reference profiles from either external single-cell RNA sequencing (scRNA-seq) dataset or query data itself (see **Methods** section), the primary requirement is that they accurately represent the expressional centroids of the major cell types present in the query dataset. Capturing fine cell types, on the other hand, is less critical for FastReseg, as the differentiation between closely related subtypes often offers limited resolution in transcript-based local evidence and thus is of minimal utility in detecting and correcting segmentation errors.

These “transcript scores” form the core operating mechanism of FastReseg and quantify how well each transcript’s expression profile matches the expected profile of its designated cell type. This involves assessing the consistency of transcripts within a neighborhood defined by initial image-based cell segmentation—ensuring they align in terms of log-likelihood ratios against the same cell type—and examining their spatial distribution. Such scores not only represent the goodness-of-fit of the transcript to the cell type but also highlight any spatially dependent patterns that may indicate the presence of contaminating source cell and thus segmentation inaccuracies (**Figure 1**C). This dual approach ensures that both expression and physical space are considered, enhancing the accuracy of detecting and correcting segmentation errors.

### FastReseg Framework

FastReseg’s workflow (**Figure 1**D) is a three-tiered process which begins with two essential inputs: the initial results of cell segmentation, typically derived from morphology images and including the transcript level of cell assignment in space, and a matrix detailing transcript scores across all genes and cell types (**Figure 2**B). The first tier of the process involves evaluating each cell’s transcriptional spatial pattern with a “spatial doublet test,” where each cell’s transcript scores under its most probable cell type given its overall single-cell expression profiles are analyzed to identify any evidence of spatial dependency. Cells exhibiting an enrichment of poorly fitting transcripts in specific areas are flagged for potential segmentation errors (**Figure 2**C). This test, which can be performed very rapidly and massively in parallel, lets us redact cells with no evidence of segmentation error from more computationally intensive downstream operations. The second tier of the process focuses on cells flagged from the spatial doublet test and attempts to segregate high-score transcripts (well-aligned with the cell’s type) from low-score transcripts (poor fits) via a spatial model trained for each flagged cell (**Figure 3**B). The regions rich in low-score transcripts are considered likely candidates for contaminating transcripts due to segmentation errors. Depending on the analytical needs, these erroneous regions can either be excised to clean up the dataset or subjected to further refinement steps in third tier. For cells with identified misassigned transcript groups, FastReseg applies a series of heuristics to determine the appropriate corrective action (**Figure 4**): whether to trim these areas from the dataset, merge them into an existing neighboring cell, or establish them as new, independent cells. This decision-making process is guided by the spatial and expression context of the transcript groups, ensuring that corrections respect both the physical and expressional landscapes of the tissue. By integrating sophisticated transcript scoring with practical segmentation refinement in a modular pipeline, FastReseg offers a powerful tool for researchers seeking to delve deeper into the cellular complexities of biological tissues.

**Figure 2.**
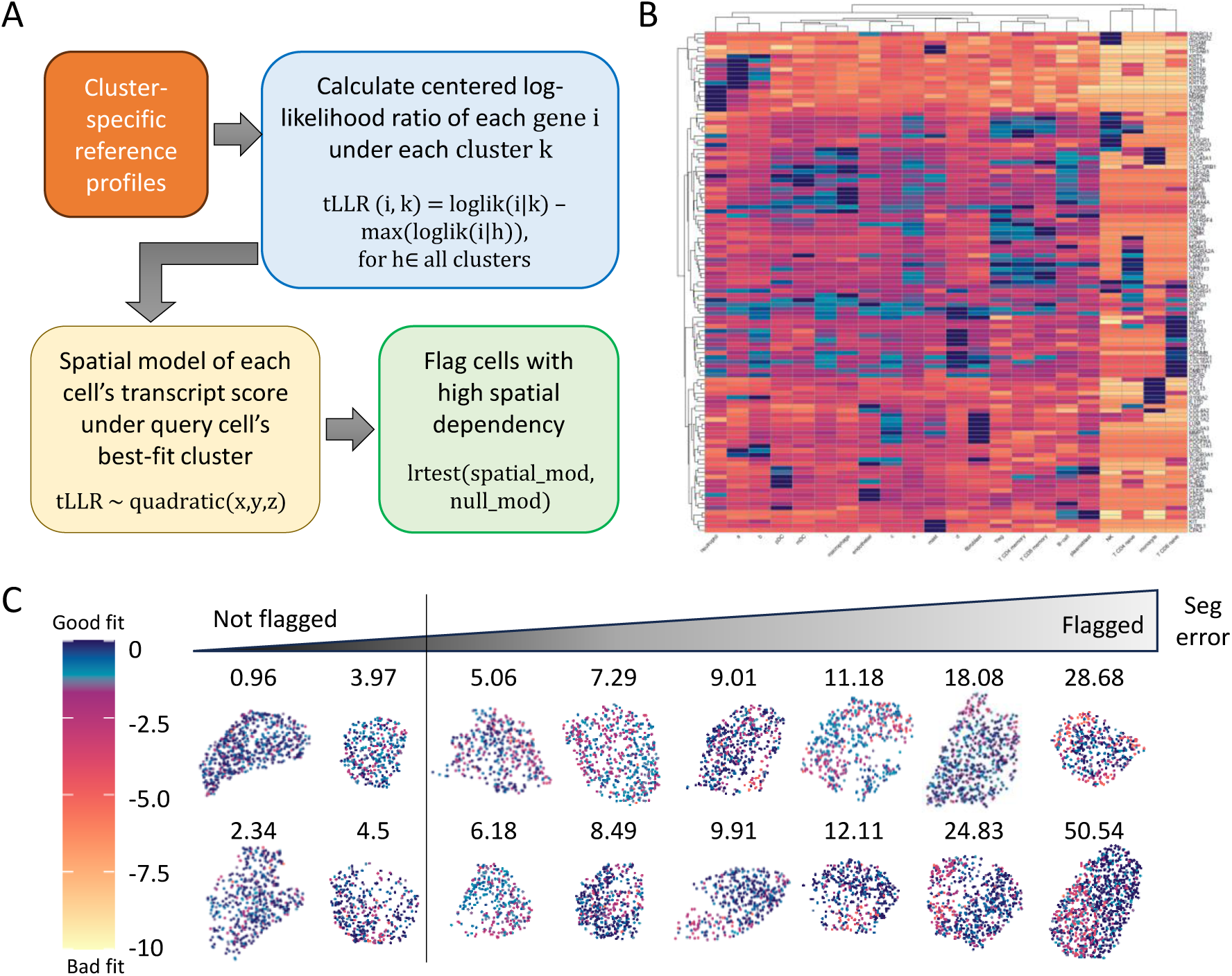
Detection of cells with putative segmentation errors based on local transcriptional profiles in both expression and physical space. (A) Flowchart of how FastReseg identifies cells with potential segmentation errors. Cluster-specific reference profiles are utilized to calculate the centered log-likelihood ratio (tLLR score) for each gene under each reference cell cluster/type, where tLLR(i, k) is derived by subtracting the maximum log likelihood of gene i across all clusters from its log likelihood under cluster k. A linear regression model is then trained to simulate the spatial pattern of each cell’s transcriptional score under its best-fit cluster given the overall expression profiles of query cell observed in current cell borders. FastReseg would further evaluate the degree of spatial dependency of the transcript score pattern of each query cell via log-likelihood ratio test between the spatial-dependent model and the invariant null model. Cells with high spatial dependency are flagged as cells with putative segmentation errors. (B) Heatmap of transcript tLLR score for top marker genes (rows) across different reference cell types (columns). The reference cluster-specific profiles (see “example_refProfiles” object inside the package) were derived from an example spatial transcriptomics dataset for melanoma tissue section whose cell types were assigned based on semi-supervised cell typing algorithm with novel clusters using “a” to “f” letters as names. The color scale, matching (C), illustrates the range from good to bad fit, highlighting the variability of score for classic marker genes in different reference cell types. (C) XY scatter plots of transcript tLLR scores within example cells with varying likelihood of segmentation error, as determined by the degree of spatial dependency (defined as −log_10_(p), and annotated on top of each cell) of their tLLR patterns under their most probable cell types. Cells with spatial dependency scores higher than the provided cutoff (default to 5, shown as the vertical line above) are flagged as improperly segmented cells.

**Figure 3.**
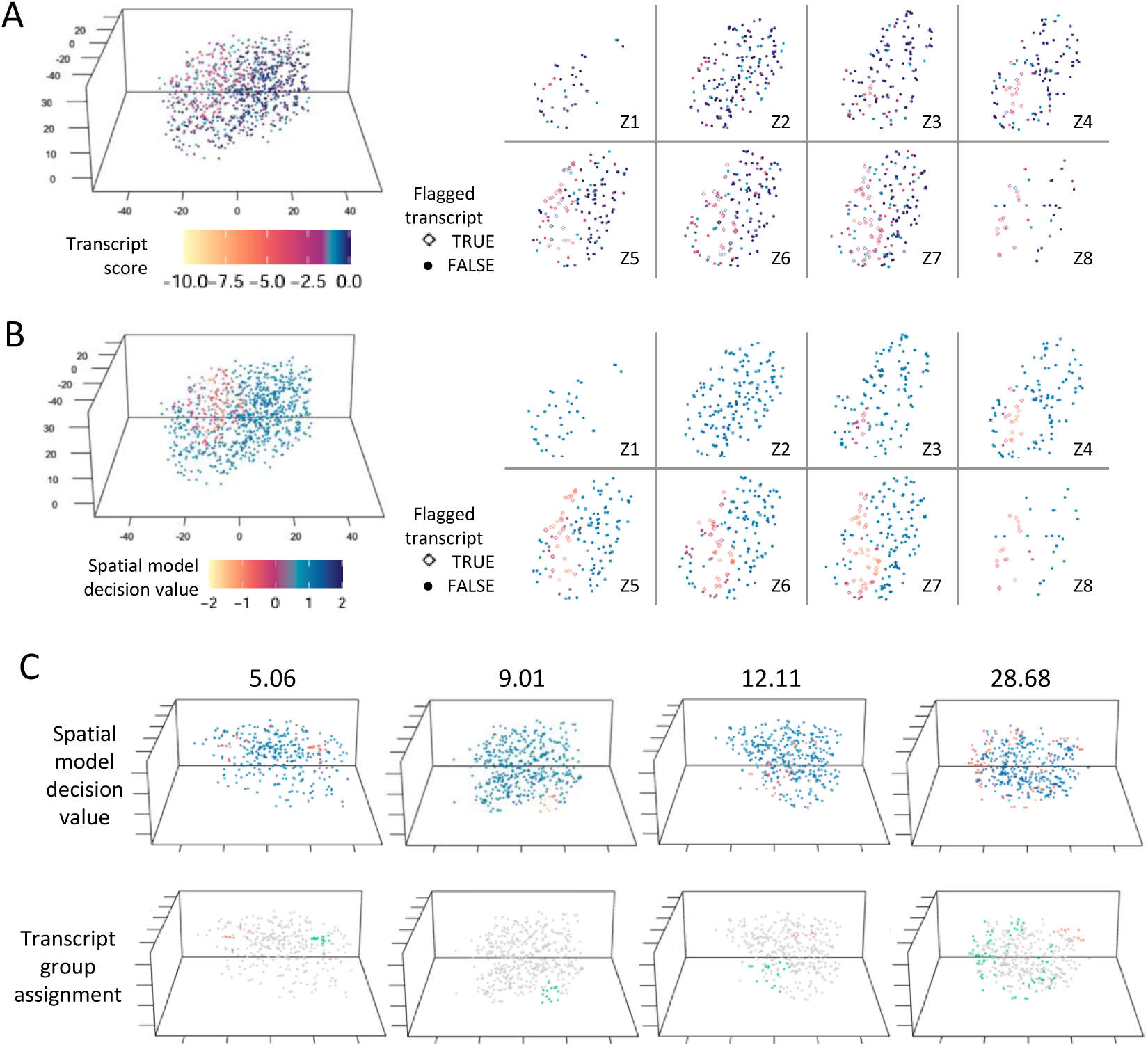
Detection and segregation of misassigned transcript groups. (A) XYZ scatter plots of transcript tLLR scores for a flagged example cell with spatial dependency score of 50.54, visualized both in 3D (left) and across multiple 2D Z-slices (right). Each point represents a transcript, colored by transcriptional score under the most probable cell type given the cell’s overall gene expression profile. This visualization highlights the spatial distribution of potentially misassigned transcripts within the cell. (B) XYZ scatter plots of the decision values produced by a Support Vector Machine (SVM) model trained to predict whether a given transcript would have transcript score below the −2 cutoff given its spatial coordinate within the same cell shown in (A). Negative decision values correspond to below-cutoff poor-fit prediction by the SVM model. In both (A) and (B), the shape of the points corresponds to the classification predicted by the SVM-based spatial modeling of the given cell. Transcripts predicted to have transcript score below cutoff are treated as flagged misassigned transcripts and depicted as hollow diamond points in the XY scatter plots for multiple Z-slices on the right of (A) and (B). The difference in the spatial pattern of transcript tLLR scores and SVM decision values of same cell provides insights into the spatial constraints on predicting misassigned transcripts. (C) Additional examples of cells with varying degrees of spatial dependency on their transcript scores (5.06, 9.01, 12.11, 28.68). For each cell, transcripts predicted as poor-fit with negative decision values by the SVM are further segregated into spatially distinct groups, shown in different colors at bottom panel. These groups indicate the likely origins of those flagged misassigned transcripts from different neighboring source cells, providing a basis for further targeted reassignment and refinement of cell segmentation.

**Figure 4.**
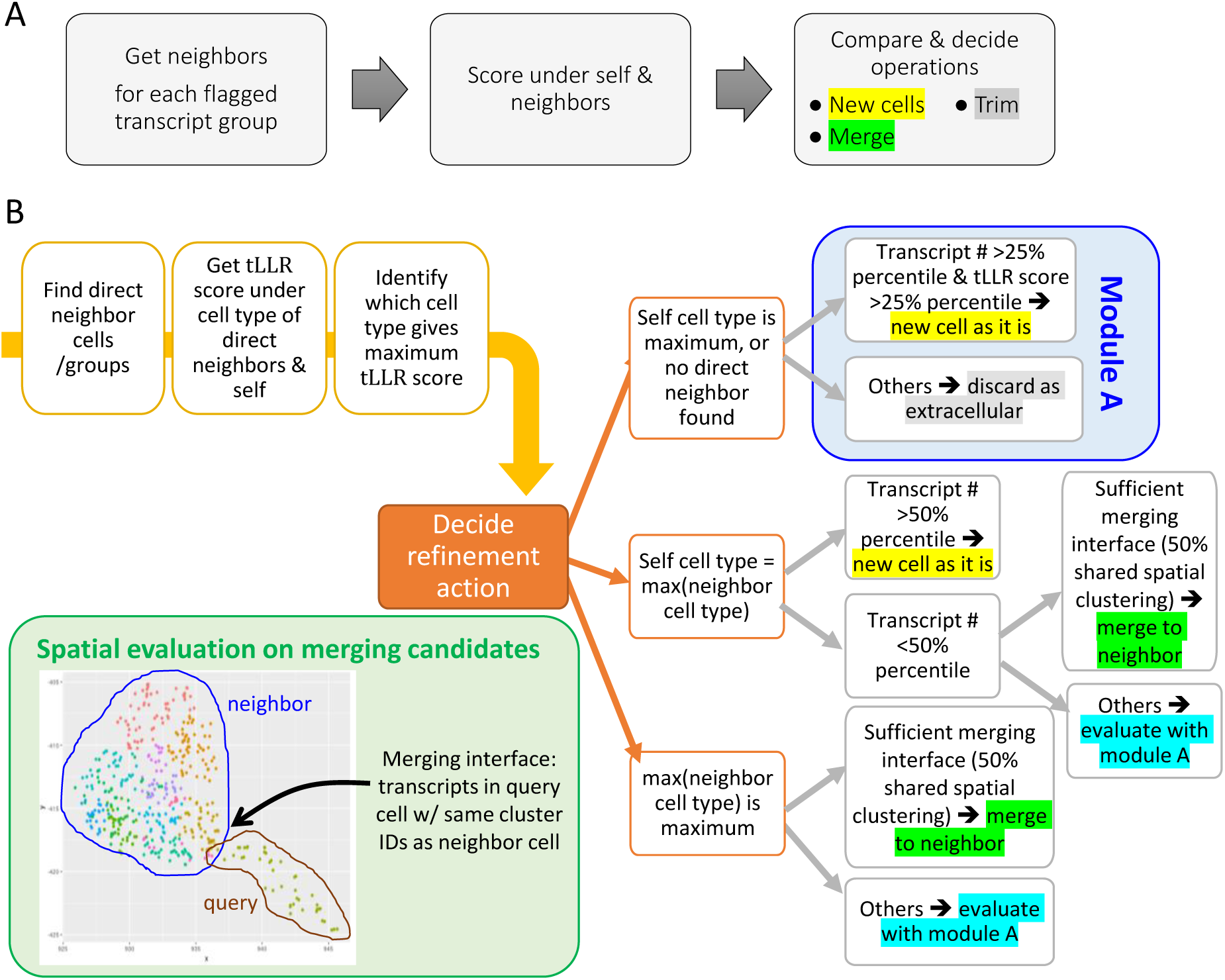
Heuristic-based approach to evaluate and reassign poorly fitted transcript groups. (A) High-level workflow for transcript reassignment, involving the identification of direct neighboring cells or groups in space, assessing total transcript tLLR scores under the most probable cell types of both the cell/group itself and its neighbors, and determining refinement actions such as creating new cells, trimming, or merging. (B) Decision tree and detailed criteria for deciding refinement actions based on the analysis of physical and expression contexts with respect to baseline distribution of transcript number and total transcript score of relevant reference cell types observed in the query dataset under the original cell segmentations. The inset plot at lower left corner in (B) shows the XY scatter plot of transcripts within an example pair of merging candidates, where Leiden clustering is performed on transcripts’ spatial coordinates to segregate them into different spatial clusters shown in different colors. The merging interface, as defined by the fraction of query transcripts sharing the same spatial clusters as the transcripts in its mering candidate partner, is used to evaluate whether sufficient physical contact exists between the pair of merging candidates, emphasizing the necessity of spatial constraint on a valid merging event.

### Spatial Doublet Test

The “spatial doublet test” in FastReseg is designed to identify cells whose expression profiles might erroneously include transcripts from adjacent cells, a situation we term “spatial doublets”. This concept extends traditional doublet detection from scRNA-seq by addressing unique challenges inherent in spatial data. Unlike scRNA-seq, where doublets involve complete merging of cellular transcripts, spatial doublets may only contain fragments of a contaminating cell’s transcriptome. Additionally, spatial transcriptomics provides the advantage of precise transcript locations, allowing for the utilization of spatial information beyond the capabilities of traditional scRNA-seq’s “bag of RNA” approach.

Among various techniques appropriate for detecting spatial doublets, FastReseg uses polynomial regression to efficiently tackle this issue, making the assessment of millions of cells amenable. Specifically, FastReseg employs a quadratic model that predicts transcript scores based on the three-dimensional positions of transcripts within a cell (**Figure 2**A). For each cell, the algorithm compares the fit of this quadratic model to a null model, where the position is assumed not to influence transcript score. The p-values of likelihood ratio test between these two models serve as a metric to evaluate the presence of spatial doublets. Cells exhibiting small p-value suggest significant spatial dependencies in their transcript distributions and are flagged as potentially suffering from segmentation errors (**Figure 2**C). This approach not only highlights the cells with likely errors but also effectively filters out those with no apparent issues. With a reasonable cutoff (*p* ≤ 10^−3^ *or* 10^−5^), FastReseg excludes 80-90% of cells with no strong evidence of segmentation errors from further analysis (**Figure 5**A), substantially reducing unnecessary computational workload in subsequent steps of the pipeline.

**Figure 5.**
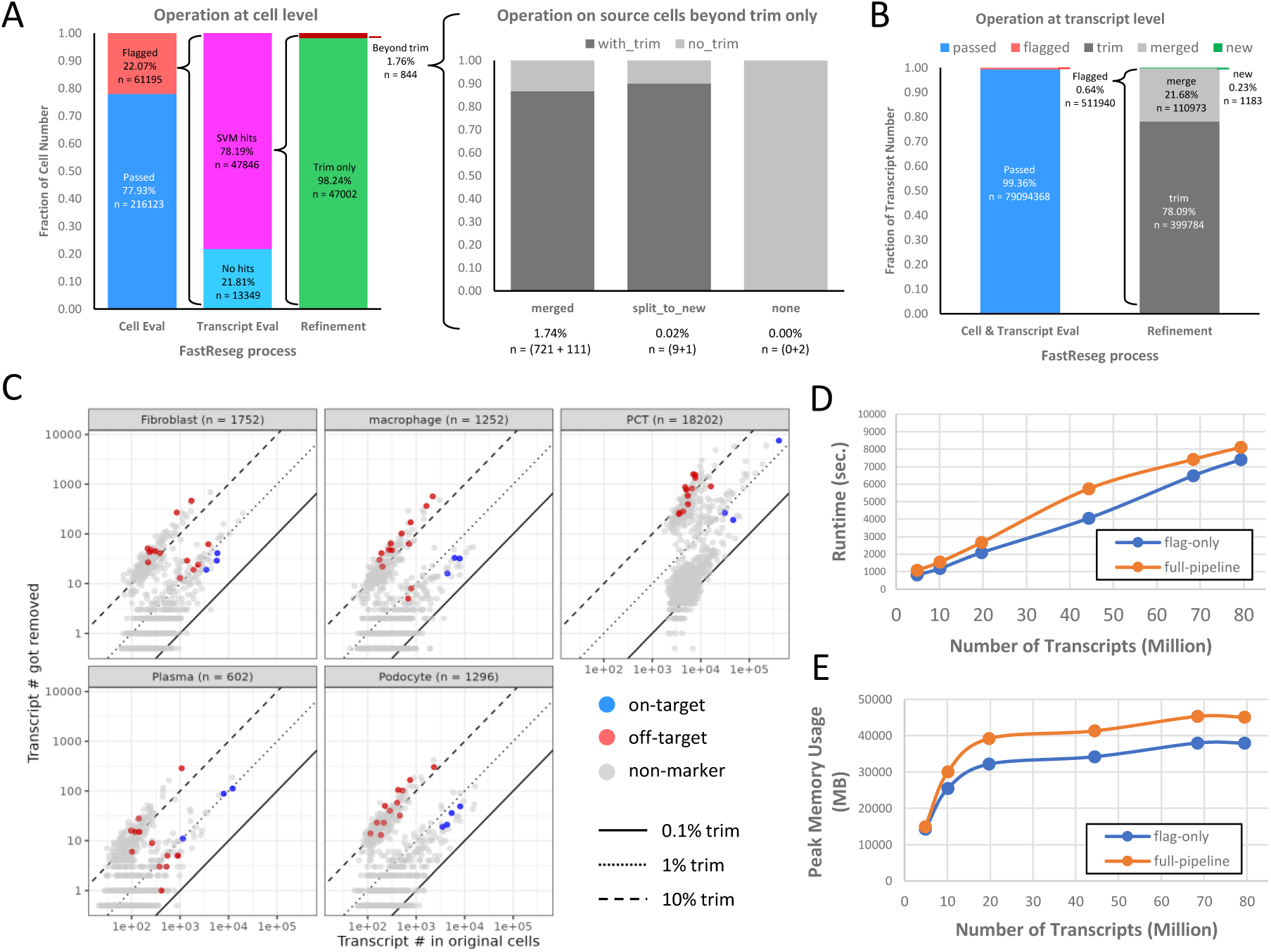
Performance evaluation of FastReseg on an example kidney dataset. (A) Bar plots on the composition of actions taken at cell level throughout the FastReseg workflow. While majority of cells (98.24%) with misassigned transcript groups identified by SVM modeling have received trimming during the transcript refinement stage, the bar plot at the center highlights the different refinement actions applied to the original host cells on top of trimming some transcripts to extracellular space, including merging those misassigned transcripts to neighboring cells (“merged”), designating them as new cells (“split_to_new”), or returning back to the host cell (“none”, n = 2 cells). Color legend of “with_trim” and “no_trim” indicates whether the host cells received more than one refinement actions. (B) Bar plots on the breakdown of operations conducted at the transcript level, illustrating the extent of transcript reassignment across the dataset. (C) Scatter plots of transcript number of each gene for each cell type. Only cells that received transcript trimming are included in this before-vs-removed comparison to show the impact of FastReseg refinement on original host cells under different cell types. The cell number included in each subplot is noted in the subtitle next to the cell type of interest. Dots for each gene are colored based on whether they are on-target or off-target marker genes for the cell type of interest. Genes that are not considered as mutually exclusive markers are colored in gray as “non-marker”. See Methods section for the mutually exclusive marker genes used in this analysis. The solid, dotted and dashed lines indicate trimming at different percentages (0.1%, 1%, 10%). (D) Runtime and (E) peak memory usage of FastReseg flagging-only and full-pipeline processing on datasets containing different transcript numbers. The statistics reported are based on processing using 75% cores of an Amazon r5b.4xlarge instance (16 vCPU, 128GB memory). The input spatial datasets were subsets of the CosMx kidney dataset with different numbers of fields of view (FOVs) containing varying cell numbers in their original cell segmentation.

### Detecting Misassigned Transcripts via Support Vector Machine

In FastReseg, once a cell is flagged for potential segmentation errors via the spatial doublet test, the next crucial step involves identifying and isolating misassigned transcripts within these flagged cells. Given the fact that transcripts belong to the same source cell would stay in proximity to each other in physical space, FastReseg looks for spatially contiguous regions enriched with molecules of low transcript scores and considers transcripts residing in those regions to be the contaminants due to segmentation error irrespective to their individual transcript scores. Frameworks used in earlier works, such as hidden Markov random fields or building of molecular nearest-neighbor networks, are theoretically appealing but computationally expensive. FastReseg utilizes a more efficient strategy: a support vector machine (SVM) [11] with a radial kernel is fitted to the spatial distribution of the assigned transcript scores within each flagged cell under each cell’s most probable cell type. The trained SVM model can effectively predict low versus high transcript scores based on their 3D positions within the query cell (**Figure 3**A-B). The radial kernel is particularly suited for this task as it accommodates non-linear boundaries shaped by the physical contours of cells and can identify multiple, spatially distinct groups of low-score regions (**Figure 3**C) that may emanate from different source cells in the neighborhood of the query cell.

The threshold defining low vs. high transcript scores is a critical turning parameter of the SVM-mediated detection of misassigned transcripts. FastReseg employs a default threshold of −2, which corresponds to a p-value of approximately 0.046 under the interpretation of transcript score as a log-likelihood ratio between two competing hypotheses about its cell type origin (see **Supplementary Materials**). Adjusting this threshold higher can lead to a more aggressive approach which flags more molecules with intermediate evidence of poor fitting given the higher p-value. Moreover, FastReseg offers the flexibility to modify SVM parameters, such as the gamma and cost, for fine control on the responsiveness of misassignment detection with respect to singlet transcript molecules. While the default value of SVM parameters set by the FastReseg package offers a commendable baseline for analyzing any new spatial dataset, users are free to explore and tailor their setup to the characteristics of the dataset of interest, such as the data richness—in terms of physical molecular density and gene content diversity—and the anticipated cellular morphology, whether it be round or exhibiting elongated protrusions. The impact of these parameter adjustments on FastReseg’s performance is illustrated in **Supplementary Figure 1**, which serves as a guideline for parameter optimization.

### Heuristic-based Reassignment of Flagged Transcripts

After isolating misassigned transcripts and segregating them into spatially distinct groups, FastReseg can proceed to segmentation refinement, reassigning those flagged transcript groups to their appropriate source cells. We opted to employ heuristic rules rather than more computationally intensive, algorithmically complex models such as the Markov Random Fields (MRFs) employed by Baysor [7], or the training-required tandem deep neural networks (DNNs) utilized by JSTA [8], or the Monte Carlo based inference approach adopted by Proseg [9]. Although these sophisticated models are theoretically appealing, they often do not offer proportional benefits in the context of sparse or fragmented datasets and might give spurious results disagreeing with morphological stain because of insufficient data points. Instead, the heuristic approach is chosen to strike an optimal balance between computational efficiency and the accuracy of segmentation refinement in spatial transcriptomics datasets which frequently comprise thin tissue sections harboring many partial cells. These partial cells typically contain limited transcriptional information, reducing the utility of applying complex models. The heuristic rules guiding FastReseg’s refinement process are grounded in a mechanistic understanding of cellular biology and the realistic distribution of transcripts within and across cells (see **Methods** section). As depicted in **Figure 4**, these rules pragmatically evaluate factors such as the distribution of transcript number and score for each cell type within the query dataset, the consistency of transcript profiles post-merging, the extent of spatial connection between merging candidates, and the feasibility of trimming transcripts to extracellular spaces when evidence suggests no presence of valid new-cell or merging event. In the next section, we will discuss the segmentation refinement outcomes generated with real-world spatial transcriptomics dataset.

### Evaluation of FastReseg performance

To demonstrate FastReseg performance, we employed a spatial transcriptomics dataset published by previous study [12]. This dataset was generated from archived clinical biopsies of human kidney samples at various stages of lupus nephritis and was collected using NanoString CosMx® platform and a 960-plex RNA panel. The cell segmentation coming with this dataset was performed using pretrained Cellpose models on 2D projected multi-channel IF images of several morphological markers (DAPI, CD45, PanCK, CD20 and CD298). Thanks to the advanced machine-learning cell segmentation models, cell borders defined in the original cell segmentation results show strong agreement with the rich information encoded in the morphological images but fail to capture the 3D volumes of cells properly and suffer from border error in regions with weak or incomplete staining, an inevitable issue observed in real-world tissue samples. With these segmentation results, the original study has performed supervised cell typing against a reference expression matrix constructed based on external scRNA-seq datasets from Human Cell Atlas and other publications. While different cell clustering approaches could be used without significant impact, we leveraged the existing cell typing results for simplicity to generate the cluster-mean profiles observed in this CosMx kidney dataset and use them as reference profiles to derive transcript score for each gene under each expected cell type.

In **Figure 5**A-B, we quantify the FastReseg outcomes across its modular process at both transcript and cell level when processing the entire kidney dataset through full pipeline. In first-tier process, FastReseg’s spatial doublet test flagged 22.07% of cells from the original segmentation results due to significant spatial dependency (p ≤ 0.001) in their transcript scores patterns. In second-tier process, 78.19% of those flagged cells were further identified by SVM spatial modeling to contain localized zones enriched of low-score transcripts (“SVM hits”). These zones, containing 0.64% of total intracellular transcripts, underwent further evaluation in the third-tier process of segmentation refinement. In this final tier, the heuristic-driven decision-making process led to over 98% of cells with SVM hits having their misassigned transcripts removed to extracellular space, impacting 78.09% of flagged transcripts. Additionally, 21.68% of all misassigned molecules were merged into neighboring cells that showed consistent transcriptional profiles in expression domain, affecting 1.74% of flagged cells. Notably, 10 original cells with SVM hits gave rise to new cells, representing 0.23% of identified misassigned transcripts. A very small subset of flagged cells (n =2), despite showing SVM hits, received no refinement action; this typically occurred when the particular group of identified misassigned transcripts was best matched with its original host cell, possibly due to only marginal spatial dependency from only a handful of source molecules. This nuance emphasizes the stringency and robustness of transcript reassignment within FastReseg. Noticeably, some host cells in original segmentation could receive multiple types of refinement action (**Figure 5**A, center panel), illustrating the diverse outcomes possible with FastReseg’s refinement process. For detailed examples of the refinement actions undertaken by FastReseg across various stage of the full pipeline, refer to **Supplementary Figure 2**.

While it’s challenging to establish ground-truth segmentation for large spatial transcriptomics dataset, we sought to track the transcript reassignment for major cell types via a list of canonical mutually exclusive markers, providing insights into the accuracy of segmentation refinement. **Figure 5**C compares the number of transcripts originally observed within cell borders to the ones removed by FastReseg for different subpopulations of host cells that have received transcript reassignment. The on-target marker genes for major cell types showed a minimal trimming rate around 1%, confirming their relevance to the cells of interest. Conversely, transcripts from off-target marker genes, which are unlikely to be present in the host cells, were removed at higher rate, often exceeding 10%. The remaining genes (“non-marker”) exhibited a binary distribution in their trimming rate, as genes with high likelihood to be expressed within the cell type of interest tend to have low trimming rate within same bin as that of the canonical on-target markers. This selective behavior demonstrates the integrity of the FastReseg’s choice of transcript reassignment and helps to reduce the ambiguity of the resulting single-cell expression profiles, offering a clearer insight into the cellular biology within the sampled tissue.

### Processing efficiency and big data

The whole CosMx kidney dataset comprises more than 79 million transcript molecules across 123 fields of view (FOVs). Each FOV covers a sample area of 0.985 mm x 0.657 mm and contains 0.4 to 1.2 million transcripts, with all its transcriptional data consolidated into a single file per FOV. The size of this large dataset poses a significant challenge for existing transcript-dominant cell segmentation methods, whose runtime and peak memory usage typically scale linearly with transcript quantity. For context, Baysor has demonstrated a runtime of ∼4 hr and peak memory usage of 53.2 GB for processing 4.66 million transcripts on an Amazon m5a.24xlarge instance that contains 96 vCPUs and 384 GB memory [10]. Proseg, which is said to be an order of magnitude faster than Baysor, reported a ∼16.7 min runtime and 3.16 GB peak memory usage with a CosMx lung cancer dataset of ∼ 3.16 million transcripts [9].

In contrast, FastReseg is designed with maximizing computational efficiency in mind and engineered to handle big datasets that are often required for studying complex biological problems in practice. FastReseg’s workflow processes transcriptional data on a per-input-file basis, markedly reducing the memory footprint required for large dataset and facilitating effective parallelization across multiple CPU cores. For benchmarking, on an Amazon r5b.4xlarge instance (16 vCPUs, 128GB total memory) while set to use 75% available cores (*i.e.* 12 cores), FastReseg maintained a relatively constant peak memory usage at 30∼38 GB across a range of input size varying from 20 million to 79 million transcripts for flag-only processing that evaluates and separates misassigned transcripts (**Figure 5**E). The full pipeline, which includes more computationally demanding steps of segmentation refinement, showed ∼ 7 GB higher peak memory usage when compared to the flag-only pipeline for the same amount of transcript input. This relatively constant peak memory usage, which is observed when the number of per-FOV input files exceeds the number of usable CPU cores, originated from FastReseg’s per-input-file data ingestion and its architecture of parallelized computation. When input files are fewer than the usable cores (*e.g.* the 2 input datasets with 5 or 10 FOVs, respectively, in **Figure 5**D-E), FastReseg’s peak memory usage scales linearly with the transcript number. If reducing peak memory consumption is a priority, further optimizations can be achieved by splitting transcripts into smaller image tiles for fewer data per input file or altering the percentage of cores utilized by FastReseg via its ‘percentCores’ argument.

Thanks to the three-tiered modular approach of FastReseg framework that directs computational resources to the cells that mostly need segmentation correction, we observed similar runtime between flag-only and full pipeline with only 4-28 min difference depending on the number of transcripts involved. When processing a typical benchmarking load of 4.8 million transcripts, FastReseg completed the flagging in just 13.6min and the full pipeline in 17.9min, making it more than 13x faster than Baysor and on par with Proseg. At 79 million input transcripts, which are unmanageable by both Baysor and Proseg due to their prohibitive peak memory consumption, FastReseg adeptly completed the full pipeline within 135 min, maintaining a peak memory usage of only 45 GB. This blend of speed and efficiency makes FastReseg a feasible tool for processing large-scale spatial transcriptomics datasets, enabling complex biological research on big sample collections to proceed smoothly.

## Discussion

In this manuscript, we introduced a novel category of cell segmentation algorithm with FastReseg, which employs transcriptomic data to rectify apparent errors in prior image-based segmentation, rather than defining cell segmentation boundaries dominantly from transcriptional data. This methodology distinguishes itself from other transcript-based segmentation algorithms by enhancing computational efficiency and reducing the risk of analytical circularity. FastReseg’s adept integration of 3D cellular structures through transcriptional data into its analysis framework enables it to pinpoint segmentation inaccuracies that evade detection in conventional 2D imaging approaches. Since it functions to evaluate and refine the output of existing cell segmentation, FastReseg will remain relevant and valuable even as image-based segmentation techniques continue to advance.

The workflow of FastReseg consists of four distinct steps: assigning scores to transcripts with respect to their initial host cells, identifying cells with putative segmentation errors through the “spatial doublet” test, flagging misassigned transcripts within these cells, and decisively managing the fate of these transcripts. We present this methodology as a versatile and easily extensible framework, with FastReseg being its first instantiation. This modular design not only allows each component to be independently enhanced but also ensures easy integration of future advancements into the pipeline. Additionally, FastReseg’s architecture excels in processing large datasets, a common challenge in transcript-based segmentation. Its ability to manage extensive data efficiently without overwhelming computational resources highlights its suitability for large-scale biological studies requiring swift and precise analysis.

The core of FastReseg’s methodology involves scoring each transcript for its fit within its respective cell, effectively transforming the discrete and computationally cumbersome data of gene names into continuous numeric scores. This simplification facilitates broader applicability across diverse gene panels and enables fast detection of cell segmentation errors with minimal computation overhead when coupled with the spatial doublet test. We anticipate this scoring system to be a valuable tool for other applications in transcript-driven segmentation analysis, facilitating broader research and development in the field.

The flagging mechanism for misassigned molecules in FastReseg is designed to be conservative, identifying transcripts that not only exhibit poor fit based on their gene identities but also align their spatial distribution with the physical shape of their putative source cells. This dual constraint approach— requiring both spatial and transcriptomic discrepancies for flagging—significantly reduces the likelihood of circular analysis, where the data used to correct the segmentation also biases the outcome. The current SVM model within FastReseg effectively identifies the most apparent errors, which improves single-cell expression profiles. By modifying parameters such as the gamma and cost in the SVM model, users can tailor the balance between spatial constraints and transcriptional distinctiveness to suit specific dataset characteristics or research needs. However, developing a more sophisticated model that can discern subtler signs of misassignment could further improve this process. Enhancing the model’s ability to separate misassigned transcripts with finer granularity would represent a significant future advancement for the pipeline, ensuring even more precise segmentation corrections.

In datasets derived from thin tissue sections, partial cells are a common occurrence. Removing all flagged misassigned transcripts following FastReseg’s criteria generally results in cleaner and more reliable data for downstream analyses, reducing the needs for extensive computational resources to segment cells with limited transcriptional information. For scenario requiring more complex segmentation refinement, FastReseg employs a heuristic approach to rapidly assign flagged transcripts to cells. This process operates under the assumption that the initial image-based segmentation provides a high-confidence identification of cells, despite ambiguity in exact cell borders and occasional errors in regions with insufficient morphological information due to poor staining or limited optical resolution in 3D. FastReseg’s rule-based correction method serves to complement and enhance the baseline segmentation results. However, its effectiveness may falter if the foundational image-based segmentation is drastically wrong across the dataset. In such cases, one can still use FastReseg’s error detection workflow to assess existing cell segmentation results. Nevertheless, it is advisable to redo the image-based cell segmentation using advanced machine learning techniques from the rapidly evolving computational visual field [13] before applying FastReseg’s refinement process.

Because FastReseg’s workflow focuses its segmentation refinement actions exclusively on cells identified as spatial doublets, it is currently not tailored to correct minor segmentation inaccuracies that may involve only one or two misassigned transcripts. Although these minor errors generally have minimal impact on most downstream analyses despite their prevalence, they could introduce artifacts in differential expression analysis, where small trends may appear statistically significant, especially when the number of cells under analysis is limited. To address this issue directly, future developments could include algorithms designed to refine minor boundary errors between neighboring cells and integrate them as an additional step within the existing FastReseg framework.

In summary, FastReseg marks a pivotal development in the field of spatial transcriptomics, offering key innovations and advantages for the detection and correction of cell segmentation errors. Its powers in processing massive datasets efficiently, along with its modular and extensible framework, positions FastReseg as a vital tool for future research. As image-based segmentation technologies advance, FastReseg’s ability to refine segmentation outputs through the integration of transcriptional data ensures that it will continue to provide valuable insights and improvements in cell segmentation accuracy. Moving forward, the continuous enhancement of FastReseg’s capabilities, particularly in addressing minor segmentation errors, will further cement its role as a crucial asset for researchers exploring the intricate landscape of spatial transcriptomics.

## Methods

### Code and Data Availability

Source code of ‘FastReseg’ package is available at https://github.com/Nanostring-Biostats/FastReseg. The processing of example dataset was performed using default configurations unless otherwise noted. The tutorial vignette of the package (https://nanostring-biostats.github.io/FastReseg/articles/tutorial.html) could be used to generate the data for reproducing figures in this study. The package also contains a small example dataset on FFPE melanoma tissue samples collected on CosMx Spatial Molecular Imager platform. The cell segmentation of this melanoma dataset was conducted with Cellpose models and derived from tissue images with DAPI nuclear stain and immunofluorescence stains against PanCK, CD45, CD298 and DAPI. This dataset also includes semi-supervised cell typing results which were generated using ‘InSituType’ package [14] and a scRNA-seq derived SafeTME matrix published for tumor-immune deconvolution in solid tumors [15]. Six novel clusters were identified in this example dataset and have cluster names as “a” to “f” letters. We use this melanoma dataset to generate figures illustrating the FastReseg workflow. See the manual of package for more information regarding this melanoma example.

The spatial transcriptomics dataset used in **Figure 5** and **Supplementary Figure 1-2** to demonstrate the performance of FastReseg algorithm is published by previous study [12]. FFPE sections of clinical biopsy kidney samples were profiled using CosMx Spatial Molecular Imager and the 960-plex CosMx Human Universal Cell Characterization RNA panel. Cell typing and initial cell segmentation was taken as it is from previous study and used to derive reference profiles for FastReseg processing. More than 79 million transcript molecules collected from 123 fields of view (FOVs) covering 76.64 mm^2^ area are included in the kidney dataset and processed by the FastReseg flag-only and full pipelines, both of which ingest the transcriptional data by FOV for small memory footprint and efficient parallel computation.

### Reference Transcript Profile Generation

The cluster-specific reference expression profiles represent the expected expression patterns of genes for distinct cell types within the sample and could be generated either from external sources or derived from the query dataset of interest. To get reference profiles from external sources, one can use previous single-cell RNA sequencing (scRNA-seq) or spatial transcriptomics datasets of similar tissue type and disease condition. Public databases for various cell atlas programs could be a good resource for such datasets. With the external dataset of choice, one can then calculate the average expression of each cell type in that dataset to be used as reference profiles. Alternatively, one could do cell typing on query dataset based on existing initial cell segmentation using any sensible clustering strategy, such as unsupervised Leiden clustering, reference-based supervised cell typing, or embedding-based label transfer. The initial cell segmentation, which could be based on the 2D morphological images of given query dataset, often has errors on the exact cell borders but is typically sufficient to capture the general gene expression differences across major cell types present in the sample. FastReseg package could then take the cell clusters assigned to query dataset and calculate the cluster-mean profiles for all provided clusters and use that as reference profiles.

### Scoring Transcripts Within Cells

Using the reference profiles provided, FastReseg assigns every transcript an expression-based goodness-of-fit score, *tLLR* (for “transcript log likelihood ratio”). The scoring process starts by scoring each gene under each cell type. FastReseg calculates the likelihood of gene *j* given cell type *k* based on the provided cluster-mean reference profiles 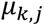:

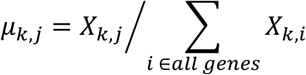

where 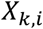 is the count of gene *i* under cell type *k* in cluster-mean reference profiles.

From these likelihood scores, we compute log-likelihood ratios for the cell type in question relative to the best-fitting cell type across all expected cell types in given query dataset:

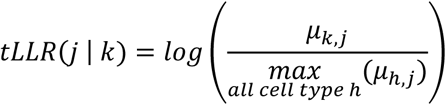

For a derivation of how tLLR scores act as a likelihood ratio test, see the **Supplementary Materials**.

Secondly, FastReseg would identify the most probable cell type for each individual cell in query dataset based on which reference cell type gives the maximum total *tLLR* score given the cell’s gene expression profile in initial cell segmentation. Then each transcript has an assigned *tLLR* score, *tLLR*(*j* | *k*) based on its gene *j* and assigned cell type *k*. This score quantifies the alignment of the transcript’s profile with the known characteristics of its most likely cell type *k*, facilitating the initial screening for potential outliers or misclassified transcripts.

### Scoring Cells for Putative Segmentation Error

Cells with segmentation errors—especially those located at the boundaries between distinct cell types— are expected to exhibit high spatial dependency in their *tLLR* transcript score profiles. This assumption could be validated by overlaying *tLLR* score profiles on membrane-stained images to confirm discrepancies between segmentation boundaries used for score calculation and the underlying biological structure. FastReseg uses linear regression model to simulate the spatial pattern of *tLLR* score profiles within a given cell and use likelihood ratio test ‘lrtest()’ from ‘lmtest’ R package [16] to compare it against a null spatial-invariant model to evaluate the degree of spatial dependency:

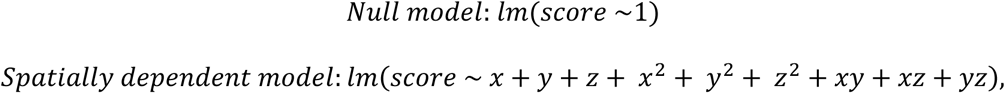

where *score* is the *tLLR* score of each transcript under the most probable cell type of the query cell and *x*, *y*, *z* are the spatial coordinates of the transcript. For transcript data with only 2D coordinates, terms associated with *z* are removed from the alternative model.

The spatial dependency score of each cell is then defined as the negative log_10_ value of the p-value from a likelihood ratio test comparing the two models. Cells with scores above a designated threshold are flagged as putative spatial doublets. Note FastReseg does not evaluate cells with very few intracellular transcripts due to low statistical power to detect spatial dependency in their transcript scores. The values used in this paper are set to 50 for minimal transcript number allowed in spatial modeling and 3 for minimal spatial dependency score of a cell flagged with putative segmentation error.

### Identification of Transcript Groups Susceptible to Segmentation Errors

To flag misassigned transcripts, FastReseg first separates transcripts within flagged cells based on the provided cutoff on *tLLR* scores into two classes and then trains a support vector machine (SVM) [11] to discern the optimal physical boundary that separates well-aligned (score above cutoff) transcripts from poorly aligned (score below cutoff) transcripts in each query cell.

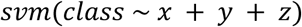

Where *z* term would be removed for 2D transcript data and additional arguments [17], like *kernal* and *gamma*, could be used to control the margin between the two classes of transcripts, adjusting the tightness of decision boundary around the hot spots of the poorly aligned transcripts (**Supplementary Figure 1**). A critical tuning parameter is the threshold for calling transcript scores as low versus high: setting this threshold higher produces a much more aggressive flagging of transcripts. FastReseg defaults to a threshold of −2, corresponding to a p-value of 0.046 under the interpretation of transcript score as a log-likelihood ratio between two hypotheses (see **Supplementary Materials** section).

The SVM-based classification approach assumes that misassigned transcripts originate from physically neighboring cells in space and thus transcripts within a physically contiguous low-score region likely belong to the same neighboring cell regardless of their individual *tLLR* scores. Therefore, transcripts predicted by SVM to have low scores are flagged as misassigned transcripts. This approach not only allows for the identification of misassigned cell-type-specific marker transcripts but also extends their impact to neighboring transcripts that may not exhibit strong cell-type-specific expression profiles. In the context of thin tissue sections, one could clean up the current single-cell expression matrix simply by trimming those transcripts, since majority of those misassigned transcripts come from the vertical overlapping of source cells with minor insertions into the assayed tissue section.

Once misassigned transcripts are identified via SVM, FastReseg could further divide them into spatially distinct groups for downstream re-segmentation process if the complete refinement of segmentation is desired. FastReseg offers two methods to segregate those misassigned transcripts in space: density-based spatial clustering (DBSCAN) and Delaunay-based spatial network analysis. The former method uses ‘dbscan’ package [18] to identify clusters of physically packed transcripts based on their local density at fast speed. The later method leverages ‘GiottoClass’ package [19] to create a Delaunay network [20] of those misassigned transcripts and identifies the transcript groups with higher accuracy based on the spatial connectivity between neighboring transcripts, facilitating the detection of potentially separate sources of misassigned molecules. The results used in this paper were generated with density-based method unless noted otherwise.

### Cell Segmentation Refinement

Each spatially distinct group of misassigned transcripts is considered as potentially originating from different neighboring source cells. To correct those putative segmentation errors, we developed a heuristic-based approach for reassigning groups of transcripts to their correct cellular origins. The process is systematically outlined in the accompanying schematic (**Figure 4**), which serves as the foundation for our segmentation refinement methodology. This section details the step-by-step process utilized in our analysis to determine the appropriate refinement action for each transcript group based on their spatial and transcriptomic context.

#### A. Defining Heuristic Thresholds

Initially, we established specific cutoffs for the number of transcripts and the transcript *tLLR* score for each reference cell type. These thresholds were determined based on the baseline distribution of these metrics across all identified cell types in the query dataset under original cell segmentation. This initial step ensures that subsequent decisions are tailored to the specific characteristics and variability inherent in the dataset for that given cell type.

#### B. Spatial and Transcriptomic Analysis

The refinement process begins with the identification of each transcript group’s direct neighbors within the tissue’s spatial context. FastReseg constructs a Delaunay network between transcript group and its neighboring cells or groups using their spatial coordinates and defines a direct neighbor based on its minimum molecule-to-molecule distance to transcripts within query group. The distance cutoff used in this paper is 2.7 µm and one can adjust this cutoff based on the observed transcript density, which could be calculated using ‘FastReseg::runPreprocess()’ function. For each group of misassigned transcripts, we assess their *tLLR* scores under the most probable cell types of itself and its direct neighbors. This dual assessment helps determine the most suitable refinement action based on local transcriptomic profiles.

#### C. Decision-Making for Transcript Reassignment

The decision tree detailed in our schematic (**Figure 4**B) directs the transcript reassignment process, employing a combination of criteria based on transcript number and total *tLLR* score under relevant cell types. Here is a clarified and concise description of this process:

- **<u>New Cell</u>**: A query transcript group is classified as a new cell if it has sufficient number of transcript molecules and a high enough total *tLLR* score under its most probable cell type. This ensures that the new cell is identified based on robust and distinct transcriptomic evidence.
- **<u>Merging</u>**: Transcript groups that do not meet the criteria for a new cell are assessed for potential merging. This assessment considers their highest *tLLR* score under the most probable cell type of their direct neighbors. A merging event is considered valid if the transcript group aligns transcriptionally with a neighboring cell or group and exhibits a substantial physical merging interface. Specifically, at least 50% of the transcripts within the query group must share the same spatial clusters as those in its adjacent merging candidate. This criterion not only confirms a strong spatial connection between the query cell and its potential merging partner but also maintains the geometric properties of the resulting merged cells. This spatial constraint for merging may be adjusted for tissue samples known to contain elongated cells or cells with protrusions, by modifying the threshold for the minimal fraction of transcripts under shared spatial clusters.
- **<u>Trimming</u>**: Transcript groups that fail to meet the criteria for either a new cell or merging are considered as extracellular. These groups typically contain only a few transcripts and are either transcriptionally distinct from all direct neighbors or lack sufficient spatial clustering overlap with any transcriptionally aligned neighbor. Such groups are likely derived from source cells that were vertically overlapping with the query cell in the original intact tissue but do not retain enough material in the thin tissue section used for the dataset.

These criteria ensure that each transcript group is evaluated based on both quantitative and spatial metrics, providing a systematic approach to accurately reassign those poorly fitted transcript groups in a context-sensitive way in spatial transcriptomics dataset.

### Evaluation of Marker Gene Distribution

We use a list of mutually exclusive marker genes of chosen cell types to evaluate the impact of FastReseg’s segmentation refinement step. Cells are grouped by their cell types in the original dataset and included for analysis if they receive trimming of misassigned transcripts. The table below shows the cell-type-specific on-target marker genes used to evaluate the kidney spatial dataset in **Figure 5**C.

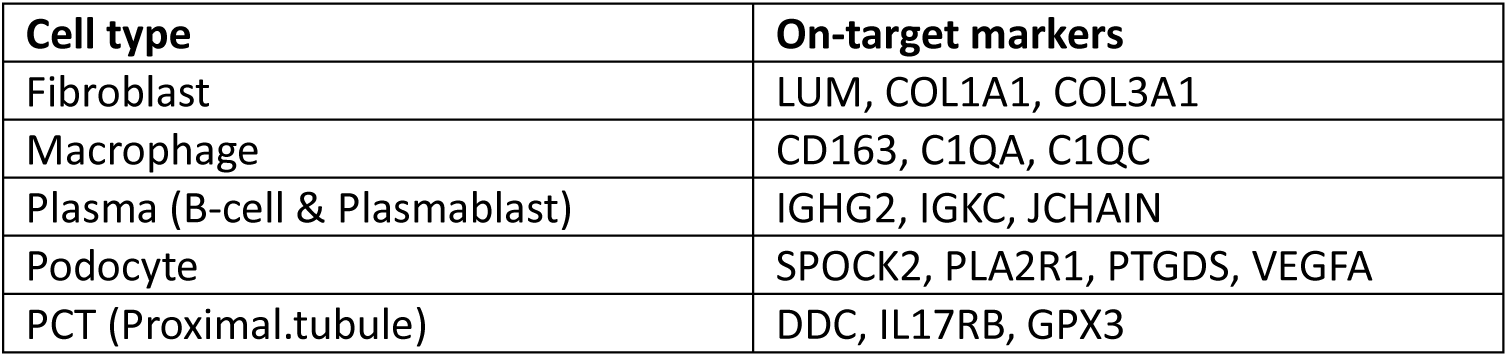

| Cell type | On-target markers |
| --- | --- |
| Fibroblast | LUM, COL1A1, COL3A1 |
| Macrophage | CD163, C1QA, C1QC |
| Plasma (B-cell & Plasmablast) | IGHG2, IGKC, JCHAIN |
| Podocyte | SPOCK2, PLA2R1, PTGDS, VEGFA |
| PCT (Proximal.tubule) | DDC, IL17RB, GPX3 |

For each cell type, on-target genes are defined as the cell-type-specific marker genes that are characteristic of that particular cell type. Conversely, off-target genes are defined as marker genes that are typically associated with other cell types and thus are the on-target genes of all other cell types in the table above. The “non-marker” category includes all the other genes present in the dataset but not included in the table above. Each subplot in **Figure 5**C displays the transcripts number of each gene within cells of the corresponding type in original dataset in x axis and the corresponding transcripts number removed by FastReseg refinement in y axis, providing a clear visual representation of the algorithm’s impact on correcting transcript misassignment in complex tissue samples.

### Characterization of Computation Efficiency

To assess the computation efficiency of FastReseg processing, we subset the CosMx kidney dataset to contain different number of fields of view (FOVs) with varying number of transcripts and cells in their original cell segmentation, as shown in table below.

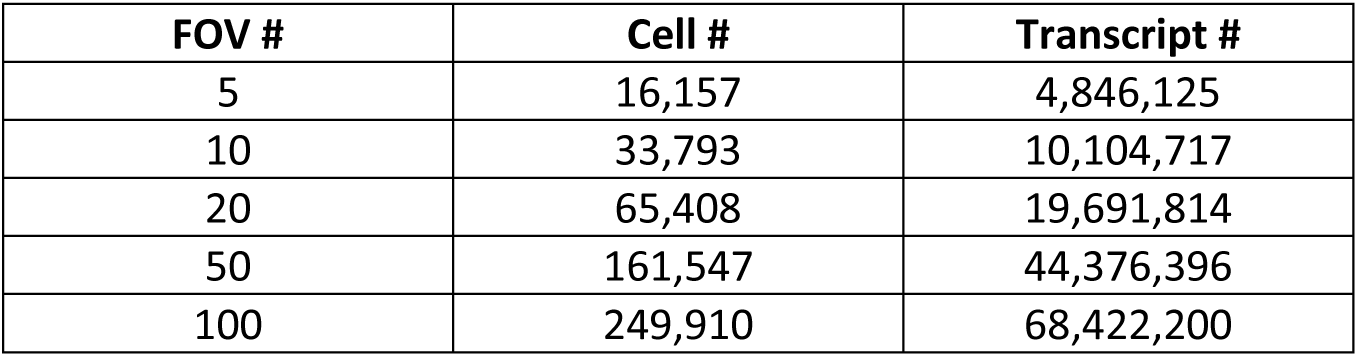
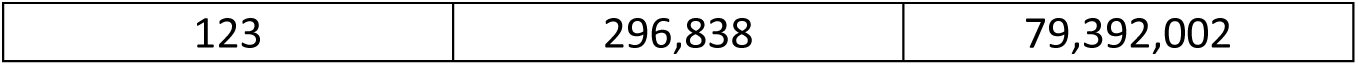

| FOV # | Cell # | Transcript # |
| --- | --- | --- |
| 5 | 16,157 | 4,846,125 |
| 10 | 33,793 | 10,104,717 |
| 20 | 65,408 | 19,691,814 |
| 50 | 161,547 | 44,376,396 |
| 100 | 249,910 | 68,422,200 |
| 123 | 296,838 | 79,392,002 |

Transcriptional data including their gene identity, 3D spatial coordinates and original cell ID assignment come as multiple files in one-file-per-FOV format from the original CosMx assay platform. We found this to be beneficial to keep memory footprint small and speed up the processing since FastReseg workflow does parallel computation at per input file level. Thus, we recommend splitting transcriptional data into image tiles for any spatial datasets collected from other platforms. We then processed those subsets through either the full pipeline or up to the point of isolating misassigned transcripts using ‘FastReseg:: fastReseg_full_pipeline()’ and ‘FastReseg:: fastReseg_flag_all_errors()’ functions, respectively. All the processing included in this study was conducted on an Amazon r5b.4xlarge instance containing 16 vCPU and 128GB memory and set to use 75% of available cores (i.e. 12 cores). The runtime was measured using ‘system.time()’ in base R. For peak memory monitoring, we wrapped the FastReseg processing in a single R script, launched it as command line and then used ‘ps aux’ command in Linux to monitor the memory usage of all processes (PIDs) associated with the R script at 10 seconds interval. The real-time memory usage of FastReseg processes was calculated as the sum of RSS (Resident Set Size, the non-swapped physical memory) across all the relevant processes and the maximum value of the real-time memory usage throughput the pipeline was then reported as the peak memory usage of the entire pipeline in **Figure 5**E.

## Supplementary Materials

### Justification of Transcript Score as a Likelihood Ratio Test

Consider a transcript inside a given cell and assume the cell’s cell type is known, designated as “cell type 0”. We employ a likelihood ratio test to quantify the evidence provided by the transcript to either support or contradict the cell type in question. The null hypothesis posits that the given transcript originates from “cell type 0”, while the alternative hypothesis considers the transcript as potentially arising from any other cell type. In gene expression domain, the only data point provided by the transcript molecule is its gene identity. The likelihood ratio between these hypotheses is then computed as follows:

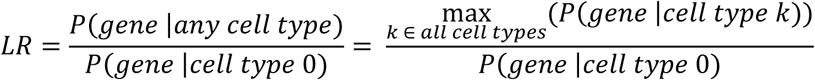

Likelihood ratio tests operate from the theory that the log-transformed likelihood ratio, −2*log*(*LR*), adheres to a chi-square distribution with the appropriate degree of freedom whose value is usually based on the number of parameters in the model. For our setup, we have only one degree of freedom given our choice of cell type that maximizes *P*(*Gene* | *cell type k*). This implies that a transcript score of −2 (the default threshold of good versus bad fits in FastReseg’s SVM step) rejects the null hypothesis with a p-value of 0.046, thereby effectively discriminating transcripts with distinct expression characteristics from the ones in agreement with their host cells’ overall expression profiles.

### Effects of SVM Configurations on Detection and Refinement of Misassigned Transcripts

To elucidate the impact of different SVM configurations on the processing dynamics of FastReseg across various stages of the workflow, a series of experiments were conducted using a kidney spatial dataset. These explorations, detailed in **Supplementary Figure 1**A-D, investigate the ramifications of altering 3 key SVM parameters: the score cutoff delineating bad-fit from good-fit transcripts, the gamma value of the SVM kernel affecting the model’s responsiveness to data complexity, and the cost parameter, which modulates the balance between achieving low error rates and maintaining a smooth decision boundary. The collective influence of these parameters determines the sensitivity and specificity of SVM modeling in identifying regions enriched with poorly fitting transcripts—areas that are potentially indicative of contamination due to segmentation error.

**Supplementary Figure 1**B reveals a direct correlation between the stringency of the score cutoff and the number of cells with successful detection of misassigned transcripts during the SVM-mediated transcript evaluation phase. Configurations with a more positive transcript score cutoff (Configurations C & D compared to A) tend to identify more cells containing “SVM hits”. These hits represent regions where misassigned transcripts are detected via SVM spatial modeling. Consequently, an increase in the number of cells containing SVM hits correlates with a greater total number of flagged misassigned transcripts, as shown in **Supplementary Figure 1**D. Similarly, a higher gamma value (Default versus Configuration A) allows bigger influence of individual below-cutoff transcripts to their spatial neighbors, flagging bigger regions and thus more transcripts as misassigned transcripts. On the other hand, a higher SVM cost value, as seen in Configuration B, imposes stringent penalties on prediction errors, leading to a more complex decision boundary that tightly conforms to the training data. This offsets the effects of a smaller gamma, thereby fine-tuning the SVM’s ability to distinguish between good-fit and bad-fit transcripts within the nuanced spatial architecture of the dataset.

Among the parameters tested, the transcript score cutoff exerts the most substantial influence on the outcome of the segmentation refinement process, as evidenced in Supplementary Figure 1C-D. More aggressive flagging of transcripts as bad fits, driven by more relaxed score cutoff or elevated gamma values, typically results in a higher proportion of transcripts and cells undergoing complex corrective actions, extending beyond mere trimming to extracellular regions. This aggressive flagging often leads to the emergence of new cell identities from those SVM-predicted bad-fit transcript groups, as they contain enough molecules to provide sufficient cumulative molecular evidence supporting the presence of new cells during refinement. Of note, it’s not recommended to use score cutoff much higher than −2, since higher score cutoff means classifying more transcripts with intermediate goodness-of-fit to their current host cells as misassigned transcripts.

These experimental findings illustrate the role of SVM parameter settings in shaping FastReseg’s behavior in detecting segmentation errors and further refinement. While the default values used by the FastReseg package provide a good starting point for any new spatial dataset, the user could adjust those parameters (e.g. gamma and cost) to fine tune the sensitivity and specificity of misassignment detection and thus tailor its behavior based on richness of transcriptional information (molecular density in physical space and gene content diversity) and expected cell shape (roundish versus long protrusions) observed in the dataset of interest.

**Supplementary Figure 1.**
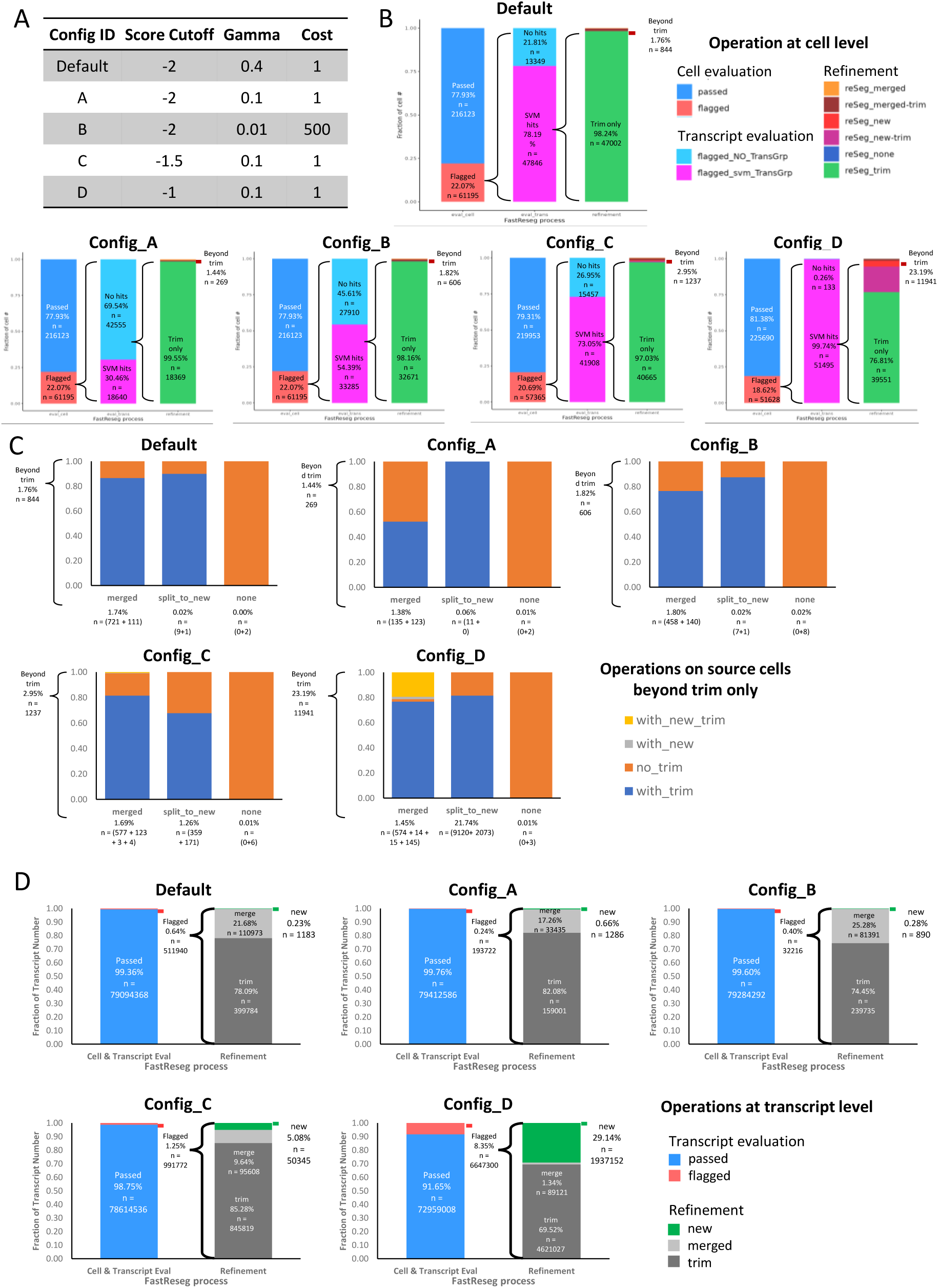
Impact of different SVM configurations on FastReseg processing of example kidney dataset. (A) SVM configurations used for performance comparison in this figure. “Score cutoff” refers to the cutoff on tLLR score to separate between well-aligned (score above cutoff) and poorly fit (score below cutoff) transcripts in SVM spatial modeling. Higher score cutoff would classify more transcripts as bellow-cutoff poor fit molecules with respect to the single-cell expression profiles of given cell. “Gamma” and “Cost” are the parameters used by ‘e1071::svm()’ function [17] to define the kernel used in training and prediction, impacting the exact shape of physical boundaries predicted by SVM spatial models. (B) Bar plots on the composition of actions taken at cell level throughout the FastReseg workflow. (C) Bar plots for composition of cell-level segmentation refinement applied to the original host cells beyond simple transcript trimming alone. The number of cells (n) included in each category is noted at the bottom of each bar. Color legend of “with_new_trim”, “with_new”, “no_trim” and “with_trim” indicates whether the host cells have received additional one or two refinement actions. (D) Bar plots on the breakdown of operations conducted at the transcript level.

### Illustrative Examples of FastReseg Processing Journey

To provide a better understanding of FastReseg’s workflow, we offer a visual narrative showcasing the distinct outcomes of its full pipeline, illustrated through examples of cells that underwent various actions during the processing of the CosMx kidney dataset. **Supplementary Figure 2** delineates different refinement scenarios into separate panels. In Panel A, we observe cells whose flagged transcripts have been allocated to form new cells. Panel C follows cells with transcripts that were merged into adjacent cells, while Panel D focuses on cells whose transcripts were trimmed to extracellular space. Each panel illustrates the comprehensive journey of cell transcripts across rows, from initial inputs to the final refinement outcomes. Additionally, **Supplementary Figure 2**B shows the single-cell distribution of spatial dependency score from the kidney dataset’s spatial doublet test results. Notably, cells that split into new cells (red line) exhibit strong spatial dependency in their assigned transcript scores. Meanwhile, cells involved in merging events (orange line) display a binomial distribution in their spatial dependency score, with a subpopulation corresponding to cells receiving transcripts from neighboring cells and showing score below the flagging cutoff (dashed black line, cutoff = 3).

The inputs of FastReseg process are depicted in rows (i) to (iii). Row (i) showcases the morphological stains that provide the visual basis for initial image-based segmentation. Row (ii) features the spatial distribution of canonical marker genes and other transcripts, highlighting the cellular composition prior to refinement. Row (iii) reveals the cell type assignments provided by the original dataset given the initial segmentation borders, setting the stage for further evaluation. Notably, cells with faint CD45 staining (red color in Row i) often have transcripts for immune-specific marker genes (Row ii). However, they are not always identified as immune cells in the cell typing results (Row iii) due to the presence of non-immune-specific marker genes. This suggests that immune cells might be present in the vertical z direction, but their cell volume may have been mostly excluded from the dataset during tissue sectioning.

The first tier of FastReseg process, presented in Row (iv) and (v), details the spatial doublet test designed to identify cells with putative segmentation errors. Row (iv) shows the spatial pattern of the transcript scores under each cell’s most probable cell type, while Row (v) reveals the spatial dependency values for each cell. Markedly, cells containing mutually exclusive marker genes and cells with boundary error between different cell types have higher spatial dependency scores and are flagged by the spatial doublet test.

The second tier of FastReseg process involves identification and segregation of the misassigned transcripts within the flagged cells. In Row (vi), the decision boundary between good and bad fits within each flagged cell is visualized by coloring each transcript according to the decision values predicted by the corresponding SVM model. Row (vii) then colors the transcripts based on their group assignments, with non-flagged transcripts shown in gray and spatially segregated misassigned transcripts in bright colors. Transcripts flagged with the same color are considered to originate from the same source cell and are evaluated as a group in the downstream refinement stage.

The culmination of FastReseg’s refinement process is captured in Row (viii) and (ix), where transcripts are depicted as dots colored according to their cell assignment before (Row ix) and after (Row ix) refinement. The 2D footprints of original cell segmentation results are represented as dim shadows beneath these transcript dots. Bright circles highlight regions where the cellular assignment of flagged transcripts has changed, demonstrating the impact of the FastReseg refinement.

Altogether, this figure serves as a comprehensive visual summary of FastReseg process, from initial segmentation and identification of potential errors to the refinement and reassignment of transcripts. Each step, visualized through detailed rows, highlights the method’s ability to detect and correct segmentation inaccuracies through a modular and systematic approach.

**Supplementary Figure 2.**
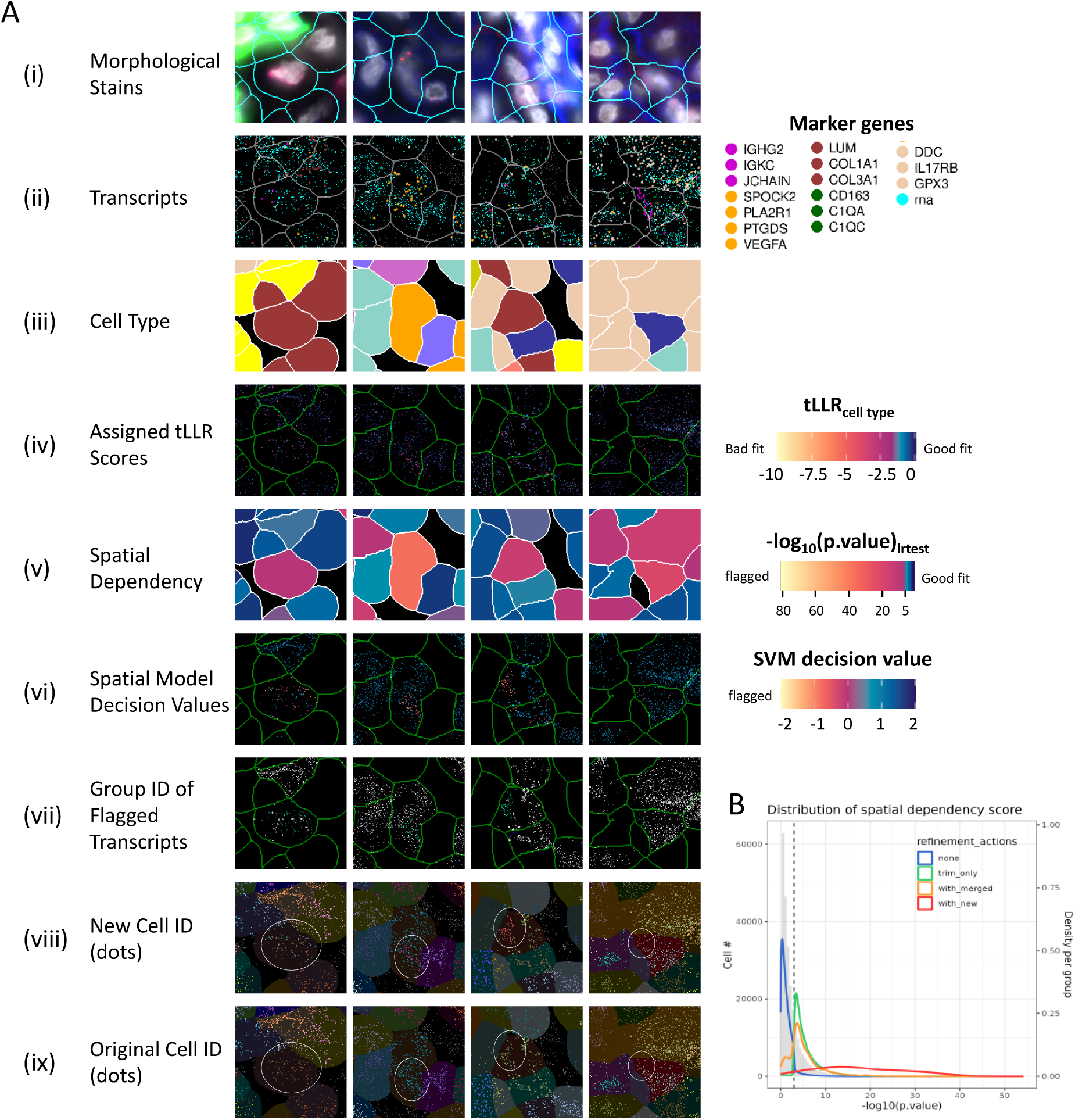
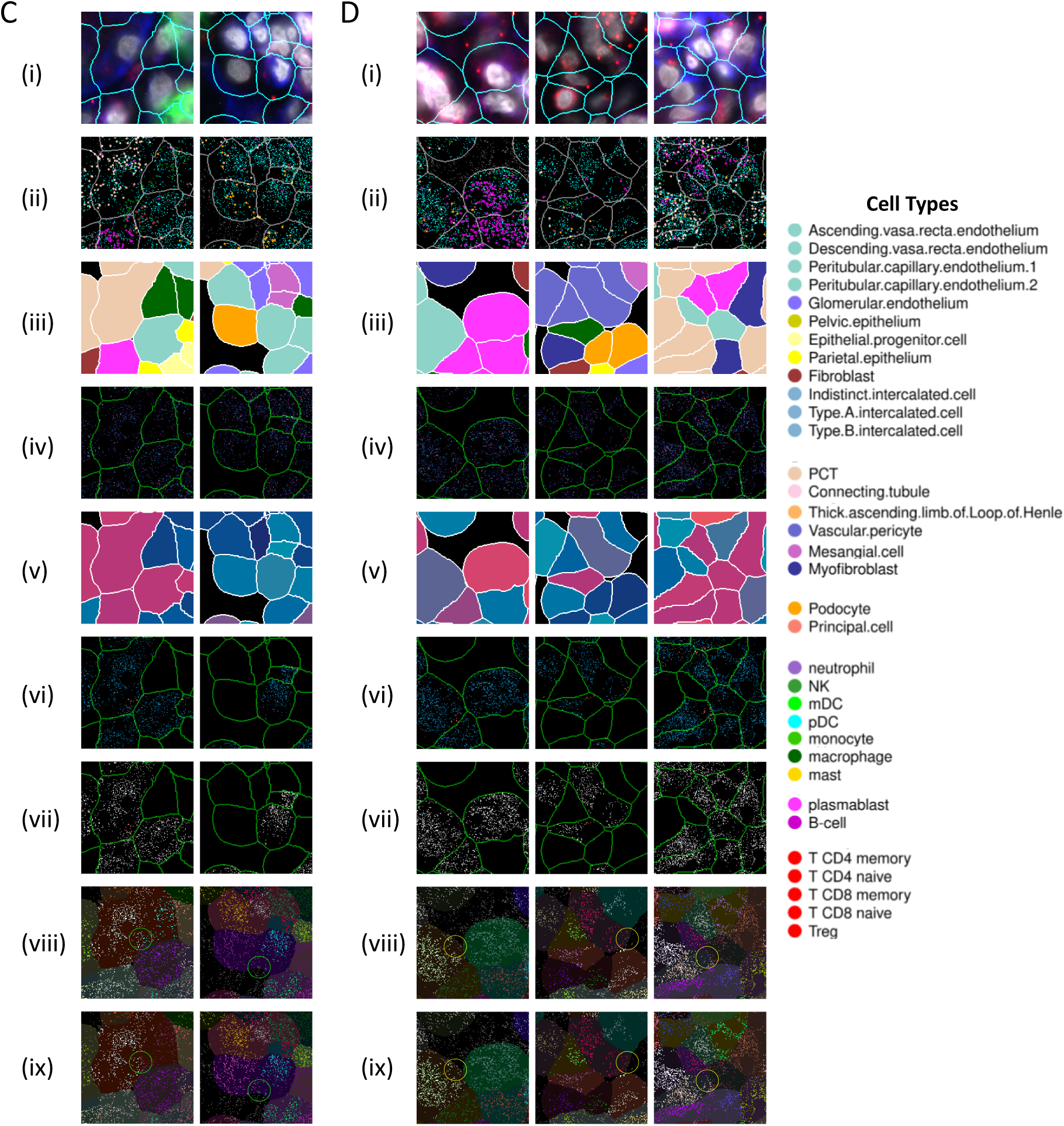
Illustrative Examples of FastReseg Process. Cells that underwent different resegment actions in FastReseg’s full pipeline processing of CosMx kidney dataset were used as demostrative examples on FastReseg’s process. (A, C-D) Panels of 2D spaital plots of various metrics, organized in rows (i-ix), for each example query cell along with its 300 px X 300 px neighborhood (54 µm dimension). Color legends are shared among Panel A, C and D, representing cells forming new cells, cells invovled in merging, and cells with misassigned transcripts removed to extracellular space, respectively. Each row represents a sequential step of the FastReseg process. Inputs for FastReseg include (i) morphological stains of query samples (PanCK: green, CD298: blue, DAPI: gray, initial cell-segments: cyan), (ii) spaital coordinates of transcripts from different canonical marker genes (colored by cell types) and other non-marker genes (cyan dots), (iii) cell typing results from original dataset using initial image-based cell segmentation results (cell area colored by cell types). The detection of segmentation errors at cell level is shown in (iv) spatial distribution of assigned transcript *tLLR* scores under the most probable cell type of each soure cell, and (v) the spatial dependency score derived from the spaital doublet test. The detection of segmentation errors at transcript level is visualized with (vi) the decistion value of good-versus bad-fit predictions from each flagged cells’ SVM spatial modeling, and (vii) the transcript group ID generated by physcial segregation of flagged misassigned transcripts (colored by group ID, with non-flagged transcripts in white, groups of flagged transcripts in different bright colors). To visualize the impact of FastReseg’s refinment actions, each transcript was colored based on their cellular assignment post-FastReseg refinement (viii) and in their original dattaset (ix), in separate rows. For easier tracking of each transcript’s location visually, the 2D footprints of original cell segmenation results are shown as dim shadows underneath those transcript dots. Regions where the cellular assignment of flagged transcripts has changed are highlighted by bright circles, illustrating context-dependent refinement actions applied to different panels of cells. Panel (B) shows the histogram (gray bars, y-axis on left) and density curves (colorful lines, y-axis on right) of single-cell spatial dependency scores observed in the CosMx kidney dataset during the spatial doublet test. Cells are grouped by their final refinement actions when calculating the group-specific density curve on their spaital dependency. The flagging cutoff used to identify cells with putative segmentation error is marked as a black vertical dashed line at −log_10_(p. value) = 3.

## Notes

### Competing Interest Statement

All authors are employees of Bruker Spatial Biology, Inc. J.M.B. also is an shareholder of Bruker Spatial Biology, Inc.

